# On the genetic architecture of rapidly adapting and convergent life history traits in guppies

**DOI:** 10.1101/2021.03.18.435980

**Authors:** James R Whiting, Josephine R Paris, Paul J Parsons, Sophie Matthews, Yuridia Reynoso, Kimberly A. Hughes, David Reznick, Bonnie A Fraser

## Abstract

The genetic basis of traits can shape and constrain how adaptation proceeds in nature; rapid adaptation can be facilitated by polygenic traits, whereas polygenic traits may restrict re-use of the same genes in adaptation (genetic convergence). The rapidly evolving life histories of guppies in response to predation risk provide an opportunity to test this proposition. Guppies adapted to high- (HP) and low-predation (LP) environments in northern Trinidad evolve rapidly and convergently among natural populations. This system has been studied extensively at the phenotypic level, but little is known about the underlying genetic architecture. Here, we use an F2 QTL design to examine the genetic basis of seven (five female, two male) guppy life history phenotypes. We use RAD-sequencing data (16,539 SNPs) from 370 male and 267 female F2 individuals. We perform linkage mapping, estimates of genome-wide and per-chromosome heritability (multi-locus associations), and QTL mapping (single-locus associations). Our results are consistent with architectures of many-loci of small effect for male age and size at maturity and female interbrood period. Male trait associations are clustered on specific chromosomes, but female interbrood period exhibits a weak genome-wide signal suggesting a potentially highly polygenic component. Offspring weight and female size at maturity are also associated with a single significant QTL each. These results suggest rapid phenotypic evolution of guppies may be facilitated by polygenic trait architectures, but these may restrict gene-reuse across populations, in agreement with an absence of strong signatures of genetic convergence from recent population genomic analyses of wild HP-LP guppies.

## INTRODUCTION

Recent evidence that phenotypes can evolve rapidly and often with surprising repeatability (convergence) has led to a re-evaluation of our expectations surrounding adaptation in nature. Particularly, understanding the genetic architecture of traits associated with both rapid adaptation and convergence can allow for insights into how adaptive variation may be maintained and be made available to respond to sudden changes in selection. Such research is important not only due to the current global circumstances of rapid environmental change, but also to understand adaptation more generally. Quantitative traits can have architectures made up of many loci of small effect (polygenic), single loci of large effect (monogenic), or an intermediate of these (oligogenic). There are currently theoretical expectations surrounding which of these are most likely to underlie rapidly adapting (Pritchard *et al.*, 2010; Jain and Stephan, 2017b) and/or convergent phenotypes (Yeaman *et al.*, 2018) but empirical evidence is only starting to accumulate.

Polygenic traits may facilitate rapid adaptation by providing a substrate of standing genetic variation to be exploited (Jain and Stephan, 2015, 2017a; Barghi *et al.*, 2019), enabling populations to adapt to shifting fitness optima by many small changes (Jain and Stephan, 2017b). Indeed, Fisher’s fundamental theorem states that the rate change of mean fitness is equal to the amount of additive genetic variance for fitness (Fisher, 1930). Conversely, rapid adaptation of oligogenic traits is expected to occur through selective sweeps, which can result in less precise shifts across the fitness landscape or ‘overshooting’ incurring genetic load (Buffalo and Coop, 2019). On the basis of this cost, it is expected that most rapid adaptation of modest changes to trait means should occur using many loci of small effect, with the exception of instances in which sudden environmental change is so extreme as to be lethal and affect absolute fitness (Bell, 2013; Whitehead *et al.*, 2017). Recent examples of suspected polygenic bases involved in rapid adaptation include shell morphologies of *Littorina* periwinkles (Westram *et al.*, 2018), immunity phenotypes in response to myxomatosis in rabbits (Alves *et al.*, 2019), and killifish adapting to anthropogenic thermal effluent runoff (Dayan *et al.*, 2019).

Similarly, there are expectations regarding the interactions between genetic architecture and convergent evolution. Whilst convergent phenotypes can arise through non-convergent genetic routes, the likelihood of convergent phenotypes having convergent genetics is expected to vary according to trait architecture. For example, polygenic traits reduce the likelihood of evolution of the same genes by increasing redundancy in the mapping of genotype to phenotype (Yeaman *et al.*, 2018; Barghi *et al.*, 2020; Láruson *et al.*, 2020). In contrast, if genetic architectures are simple and composed of few large effect loci, reduced redundancy can funnel adaptation through repeatable genetic paths. Many of the most notable examples of genetic convergence are single loci of large effect (Stern, 2013), including the *eda* gene associated with marine-freshwater armour plate phenotypes in three-spined stickleback (Colosimo, 2005), and the *optix* gene associated with wing patterning across *Heliconius* species (Reed *et al.*, 2011).

The guppies of northern Trinidad are a model system for studying phenotypic adaptation, which has provided empirical evidence for both rapid adaptation and convergent evolution. In this system, barrier waterfalls within many rivers have created replicated downstream/high-predation (HP) and upstream/low-predation (LP) habitats. Each river contains HP- and LP-adapted guppy populations that have independently evolved convergent LP phenotypes in predator-free upstream environments. LP populations are typically longer-lived, exhibiting larger adult sizes, reduced brood size, longer time to reach maturity and longer interbrood period than their HP counterparts (Reznick, 1982; Reznick *et al.*, 2001; Torres Dowdall *et al.*, 2012). Experimental translocations of HP guppies to previously uncolonised LP environments further demonstrated that LP life history phenotypes evolve rapidly (Endler, 1980; Reznick and Bryga, 1987; Reznick *et al.*, 1990, 1997, 2019; Gordon *et al.*, 2009). Guppies raised under laboratory conditions for multiple generations continue to exhibit differences between HP and LP phenotypes, indicating these traits have a heritable genetic basis (Reznick, 1982; Torres Dowdall *et al.*, 2012). Beyond this however, and despite the wealth of knowledge regarding life history evolution in these populations, little is known about the genetic architecture of these traits.

Life history traits are typically quantitative and are commonly involved in adaptation to novel or changing environments. Previous studies exploring the genetic basis of life history traits have documented everything from highly polygenic traits, such as clutch size and egg mass in great tits (Santure *et al.*, 2013) and weight of Soay sheep (Bérénos *et al.*, 2015), to traits with single loci explaining a large proportion of phenotypic variance, such as age at maturity in atlantic salmon (*Salmo salar* L) (Ayllon *et al.*, 2015; Barson *et al.*, 2015) and other salmonids (Moghadam *et al.*, 2007; Kodama *et al.*, 2018). Life history traits can often exhibit genetic covariance, with different traits sharing aspects of their genetic architecture, which can have important implications for pleiotropic constraint during adaptation (Hall *et al.*, 2006).

There are various avenues for exploring the genetic architecture of quantitative traits. For example, single locus quantitative trait locus (QTL) analyses or Genome-Wide Association Studies (GWAS) are now commonplace. However, these approaches can inflate the prominence of single loci and their inherent bias against multi-locus models has come under scrutiny (Rockman, 2012; Slate, 2013). Indeed, multi-locus analyses have resolved some empirical inconsistencies, such as the “missing heritability crisis” (Manolio *et al.*, 2009), subsequently returning much higher estimates of heritability compared with the sum of single-locus outliers for classic quantitative traits such as human height (Yang *et al.*, 2010; Yang, Manolio, *et al.*, 2011). Extensions of these multi-locus approaches have partitioned phenotypic variance into specific chromosomes, with correlations between chromosome size (as a proxy for number of functional loci) and partitioned phenotypic variance taken as evidence for highly polygenic traits (Santure *et al.*, 2013; Bérénos *et al.*, 2015; Kemppainen and Husby, 2018a). The rising prominence of multi-locus models has started to bring empirical evidence back in line with Fisher’s prediction of the importance of many loci of small effect, often dubbed the “infinitesimal model” (Fisher, 1918; Barton *et al.*, 2017). It is therefore important to explore analyses that allow for all possible trait architectures.

Using an F2 breeding cross design, we examine the genetic basis of seven life history traits in guppies: female age (1) and size (2) at first brood, first brood size (3), interbrood period (4), average dry offspring weight in the first brood (5), and male age (6) and size (7) at maturity. Our aims are to assess the relative extents to which different facets of life history traits have significant genetic elements, and whether guppy life history traits are better explained by polygenic, oligogenic, or monogenic models. By exploring the genetic bases of these traits within a system for which rapid adaptation and convergent evolution is already well-documented, we can better understand the role of quantitative genetic architecture in these processes.

## METHODS

### Crosses

Fish were second and third generation lab-reared individuals from an HP site in the Yarra river (680415E, 1193791N) and an LP site in the Quare river (696907E, 1181003N) (Figure 1). These populations have demonstrable HP-LP life history phenotypes (Table S1), and have been studied extensively in prior work (Reznick, 1982; Reznick *et al.*, 1996, 2004, 2005). Four F2 full-sib intercrosses were performed. Two crosses were performed for each cross direction in which wild-caught LP males were crossed with wild-caught HP females and vice versa. F1s were mated within cross and F2s were phenotyped and genotyped. Grandparents were also genotyped.

**Figure 1:**
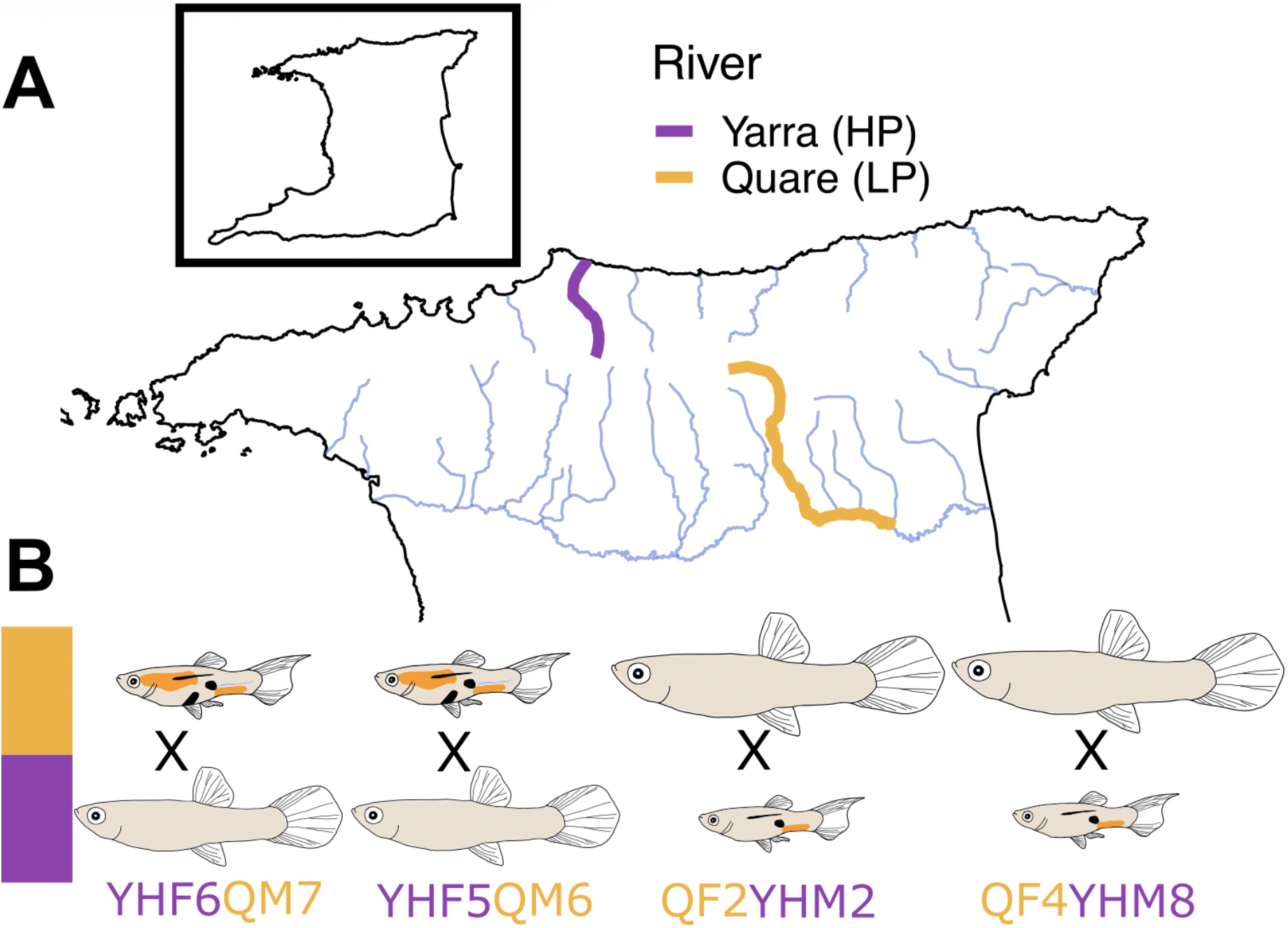
Sampling rivers in Trinidad and cross design. Sampling rivers are highlighted along with a number of major rivers from northern Trinidad’s three drainages (**A**). Four families were produced from eight grandparents (**B**), two for each cross direction. Males (smaller, colourful) and females (larger, uncoloured) in panel **B** also highlight common morphological differences between HP (smaller, less colourful) and LP individuals.

### Phenotyping and Phenotype GLMs

Life history phenotyping and rearing followed (Reznick, 1982); full rearing details are available in the supplementary materials. Size of females and males was measured under a dissecting scope with Vernier calipers following MS-222 anaesthetisation. Based on the allometric dependency of female brood size, we took residual brood size as the residual difference between observed and linear-predicted brood size based on size. Male age at maturity was judged from the development of the apical hook. Interbrood period was scored as days between first and second parturition. Offspring weight was recorded as the mean dry weight of individuals from the first litter (data from the second litter was highly correlated). Where necessary, phenotypes were log-transformed to improve fit for normality assumptions.

Rearing (mean temperature and date of birth (DOB)) and family effects on phenotypes were explored using generalized linear models (GLM) in R (v4 (R Core Team, 2020)). DOB was included as a proxy for subtle unmeasured changes in rearing conditions over time. We used backwards model selection implemented in *step()*, starting from an additive model including family, temperature, and DOB. Relevant model assumptions were checked by comparing residuals to simulated residuals in the R package *DHARMa* (Hartig, 2020). Final model term significance was determined by comparing models with and without each independent variable using *drop1()* in terms of ΔAIC and using F-tests. Where model assumptions could not be met, final model terms were taken and significance (*p* < 0.05) of Spearman’s rank correlations were used to confirm model effects. Adjusted partial R-squared was estimated for all final model variables with the R package *rsq (Zhang, 2020)* using the variance-function-based type.

### Genotyping

Genomic DNA was extracted from fin clips using an ammonium acetate extraction method (Nicholls *et al.*, 2000; Richardson *et al.*, 2001). We genotyped each individual using a RAD-seq library preparation method adapted from Poland and colleagues (Miller *et al.*, 2007; Baird *et al.*, 2008; Poland *et al.*, 2012); full genotyping details are available in the supplementary materials. Of all 661 individuals used in the final analysis, 61 were sequenced two or three times in separate libraries to account for low coverage (“merged” individuals in Table S2). To ensure optimal coverage, and reduce PCR duplicate effects, all eight grandparents were sequenced four times in four separate PCR reactions and sequencing libraries. Of the total 653 F2s, 637 (370 males, 267 females) were used for phenotype analyses due to missing phenotypes for 16 individuals.

### Bioinformatic processing

Raw read data were trimmed and adaptors removed using cutadapt (Martin, 2011). Stacks v2.5 was used for all downstream processing (Rochette *et al.*, 2019). Trimmed read data were used as input in process_radtags, with options to remove reads with uncalled bases (-c), quality filter at Q10 (-q), and rescue barcodes and RAD tags containing sequencing error (-r). Cleaned RAD tags were aligned to the male guppy reference genome using BWA-MEM (Li, 2013), and converted to bam format using samtools. Read group information was added to bam files using Picard Tools v2.6.0 AddOrReplaceReadGroups (Broad Institute, 2019) and alignments based on the same individual were merged using Picard Tools MergeSamFiles. Bam files were used as input for the gstacks module in Stacks2, using only alignments with a minimum mapping quality of 20. The final VCF contained only loci called across all individuals (-p 1), at a max-missing frequency of 80% (-r 80) and minor allele frequency (-maf) of 5%. Samples were retained with ≥ 15X average coverage, average coverage across the samples was 33.4X (Table S2) and average missing data was 0.05%. For QTL scans genotypes were imputed based on grandparental phasing (see below).

### Linkage mapping

Linkage maps were produced with Lep-MAP3 (Rastas, 2017). Pedigrees were produced for each cross by including dummy parents (one pair per cross) from which all F2s descended. Genotype likelihoods were called from the VCF input with the *ParentCall2* module, including the *-halfSibs=1* flag. Further filtering was performed with the *Filtering2* module removing markers with a MAF < 0.1 (within families) and with missing data in >10% of individuals (within families). Markers were mapped to linkage groups (LG) with *SeperateChromosomes2* modules based on a logarithm of the odds (LOD) score of 20, using all informative markers, and grandparental phase information. LGs with fewer than 20 markers were discarded, leaving 21 LGs. The two largest LGs were separated by further iterations of *SeparateChromosomes2* run over these specific LGs with an elevated LOD limit of 30. This produced 23 LGs in agreement with the guppy genome. Unmapped markers were joined to the 23 LGs with the *JoinSingles2All* module, with an LOD limit of 5. The module was iterated until no further markers could be mapped. In total, 7,256 markers of 16,539 were mapped to LGs. The module *OrderMarkers2* was then run over each LG independently to order and place markers within LGs. An initial 10 iterations were performed, with order determined by maximum likelihood. For chromosome 12, male recombination was not permitted given previous evidence that males do not recombine over the sex chromosome (Charlesworth, Zhang, *et al.*, 2020). LOD scores from these maps were used to further filter markers on the basis of support for multiple mapping within a LG (multiple LOD peaks) or if maximum LOD was within one standard deviation of the mean. These markers were blacklisted for the final *OrderMarkers2* run, in which the *evaluateOrder* flag was run over the earlier maximum-likelihood based map. Final maps were sex-averaged and trimmed according to graphical evaluation. Grandparental-phased genotypes were exported for QTL analysis. Effects of female-biased heterochiasmy (Bergero *et al.*, 2019) on linkage maps are discussed in the supplementary material.

### Heritability and multi-locus estimates of trait architecture

We used Genome-wide Complex Trait Analysis (GCTA) (Yang, Lee, *et al.*, 2011) to estimate phenotype heritability. This approach estimates the heritability of phenotypes by partitioning phenotypic variance into genetic variance (specifically of the SNPs sequenced rather than heritability in the traditional sense), random genetic effects and residual variance using the restricted maximum likelihood (REML) method within a linear mixed model. Empirical support suggests that heritability estimates using this method are comparable to those estimated from true pedigree studies (Stanton-Geddes *et al.*, 2013; Duntsch *et al.*, 2020). We first separated SNP data into males and females and, for each, estimated a genetic relatedness matrix (GRM) using all SNPs. SNPs from scaffolds were merged onto the beginning/end of chromosomes according to the linkage map (Table S3) to improve the accuracy of per-chromosome estimates. To account for genetic variance among the four families, we included the first three eigenvectors from PCA (the minimum number required to separate four families) as quantitative covariates. Specifically, these analyses are assessing within-family associations between genetic covariance and phenotypic variance, and allowing for among-family intercepts to vary. Rearing covariates were included in a trait-specific manner if these were associated with the phenotype (Table S4). To assess within-family effects, we included an additional GRM calculated with the *--make-bK 0.05* parameter. The addition of this GRM allows us to partition variance into that associated with sequenced SNPs (G1) and within-family structure (G2). At the per-chromosome level, including this additional GRM prevented model convergence in some cases (11/138 chromosome-phenotype pairs), but we observed negligible differences in heritability estimated with and without this additional GRM at the whole-genome level.

To estimate the heritability associated with specific chromosomes (*h^2^c*), we took two approaches: 1) We partitioned phenotypic variance using a model with a GRM derived from the focal chromosome and quantitative covariates; 2) We used a likelihood-ratio test (LRT) approach following (Santure *et al.*, 2013, 2015; Duntsch *et al.*, 2020) in which we compared a full model *a* (GRM based on all chromosomes except focal chromosome + focal chromosome GRM) to a reduced model *b* (without focal chromosome GRM). Quantitative covariates were included in both models. Models were compared with a LRT according to (1), with p-values taken from the chi-squared distribution with one degree of freedom.

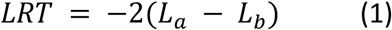

Correlations can reveal polygenic architectures. Positive correlations suggest traits are associated with many loci of small effect, assuming chromosome size is a proxy for functional loci count. Previous work has suggested p-values derived from both chromosome partitioning approaches highlighted above are comparable (Kemppainen and Husby, 2018a), so we used *h^2^c* estimations from focal chromosome GRMs (approach 1) as issues with model convergence prevented estimations of *h^2^c* on chromosomes that accounted for little phenotypic variance under approach 2 (see Table S5). Regressions were performed using the *HC_Correction()* function presented by Kemppainen and Husby (2018b), which corrects for heteroscedasticity among chromosomes and the constraint of GCTA models that prevents negative estimations of *h^2^c*.

### QTL scans

We produced two datasets for QTL-scanning. The first included fully-informative SNPs for both founding populations, i.e. SNPs that were homozygous in all eight grandparents and fixed for alternative variants between HP and LP-derived grandparents. This dataset included 1,220 SNPs, and allowed for analysis of all individuals together to increase biological power, where individuals could inherit an HP (H) or LP (L) allele. The second dataset comprised four separate datasets, one for each family in which family-informative SNPs were included for each family (homozygous, alternative, SNPs in each grandparent within families). Numbers of family-informative SNPs for each family were similar (3,436; 3,400; 3,476; and 3,476 for QF2YHM2, QF4YHM8, YHF5QM6, and YHF6QM7 respectively). These datasets provided weaker biological power, but increased resolution within families, and also allowed us to examine alleles that may have only been captured and are segregating in single crosses. It is important to note, these latter datasets cannot be used to assess loci that are Y-linked. Because all males within a family carry the same Y, the effect of different Y loci among families cannot be separated from general genome-wide relatedness within and among families that captures many autosomal loci.

We first performed single locus scans using the *scan1()* function in R/qtl2 (Broman *et al.*, 2019). We inserted pseudomarkers using *insert_pseudomarkers()* with step=1, calculated genotype probabilities with *calc_genoprob()* and an error_prob=0.002, and converted genotype probabilities to allele probabilities using *genoprob_to_alleleprob()*. We calculated a grid with *calc_grid()*, subsetted genotype probabilities to grid pseudomarkers with *probs_to_grid()*, and used this to calculate a kinship matrix with *calc_kinship()* according to the Leave-One-Chromosome-Out method (LOCO). Rearing covariates and a binary family assignment matrix were included as additive covariates. Significance of LOD peaks was determined by 1000 permutations for all models with *operm()* at an α of 0.05. For scans of the sex-determining region, inputs were merged for males and females, and the same methodology was used, with the exceptions that sex was modelled as a binary phenotype, and rearing covariates were excluded from the covariate matrix.

We also explored QTL scans that allow for multiple QTLs using the *stepwiseqtl()* function in R/qtl (Broman *et al.*, 2003) which is not available in R/qtl2. This approach assesses interactions among all pairs of loci, allowing for epistatic effects to be examined. We allowed for models with a maximum of six loci, used the imputation method, and allowed for only additive interactions among loci. Significance of LOD peaks was determined based on 1000 permutations.

## RESULTS

### Phenotypes

Both male age (GLM: *F*_3,340_ = 14.75, *p* = 4.82e^−9^) and size at maturity (GLM: *F*_3,340_ = 16.02, *p* = 9.30e^−10^) of F2s varied significantly between the four cross families (Figure S1A; Table S4). For age at maturity, cross YHF5QM6 F2s tended to mature later than all other crosses (Tukey *p* < 0.05), and for size at maturity, cross QF4YHM8 F2s matured at a larger size. Male age and size at maturity were not strongly associated with one another (correlation of individuals, Spearman’s ρ = −0.07, *p* = 0.184), but a GLM of male size at maturity by an interaction between age at maturity and family revealed a significant effect (GLM: *F*_3,340_ = 2.98, *p* = 0.031). However, within this model relationships between male age and size at maturity were both positive and negative depending on family. Rearing conditions affected both male phenotypes. Males matured earlier under increased temperatures (GLM: *F*_1,340_ = 13.75, *p* = 2.40e^−4^) and if born later in the experiment (GLM: *F*_1,340_ = 23.26, *p* = 2.15e^−6^). Larger males at maturity tended to be born later in the experiment (GLM: *F*_1,340_ = 41.51, *p* = 4.06e^−10^), but temperature did not affect size at maturity. Family status explained 10.4% and 11.7% of phenotypic variance for age and size at maturity respectively, which will not be captured by our subsequent mapping approaches.

Female phenotypes were generally less variable among families, only female size at first brood (GLM: *F*_1,265_ = 11.88, *p* = 6.04e^−4^) and offspring weight (GLM: *F*_1,249_ = 3.03, *p* = 0.030) differed significantly between families (Figure S1B; Table S4), with the latter effect only marginally significant. Consistent with a general life history axis, covariance among female phenotypes was generally high (Table S6). All female traits loaded positively onto PC1 (37.6%), with female age (loading = 0.68) and size (loading = 0.58) loading particularly strongly. PC2 (27.0%) explained residual variance associated with brood traits, with first brood size (−0.62) and interbrood (−0.59) loading negatively, and offspring weight loading positively (0.44). PC2 therefore summarises variation among females with few, heavier offspring and short interbrood periods, and vice versa. Similar to males, females reached maturity and produced their first brood earlier under increased temperatures (GLM: *F*_1,265_ = 21.65, *p* = 5.19e^−5^). Increased temperature also reduced interbrood period (GLM: *F*_1,265_ = 24.32, *p* = 1.45e^−6^) and females born later in the experiment had longer interbrood periods (GLM: *F*_1,265_ = 13.47, *p* = 2.94e^−4^). Other female phenotypes were not associated with rearing conditions (Table S4). In contrast to male phenotypes, family status explained much less phenotypic variance: 6.7% for first brood size, 5% for size at first brood, and 2.2% for offspring weight (‘family’ was dropped from other models due to low explanatory power).

### Linkage mapping

The final linkage map consisted of 6,765 markers and a length of 1,673.8 cM. There was overall good concordance between this genetic map and the recently updated reference genome (Fraser *et al.*, 2020), with the additional placement of unplaced scaffolds on all linkage groups. There were also minor structural rearrangements and inversions (Figure S2), for which corroborative support could be found from previously published HiC data (Fraser *et al.*, 2020). Typically, unplaced scaffolds were joined to either chromosome end (Table S3; Figure S2).

### Heritability and multi-locus estimates of trait architecture

Heritability varied between traits, but in almost all cases (excluding first brood size) the variance explained by within-family structure (V_G2_) was greater than the variance explained by the specific SNPs themselves (V_G1_). This is expected given RAD-sequencing is designed to capture SNPs in linkage with causal variants, rather than causal variants themselves. Estimates of heritability were greatest for male size at maturity (43.4%), offspring weight (33.4%), male age at maturity (31.3%), and interbrood period (30.2%). Estimates for the remaining female life history traits were lower, not exceeding 9.7% (female size at first brood), with standard errors that overlapped 0 (Table 1).

**Table 1:** Estimates of genome-wide heritability for each phenotype based on GCTA-GREML. Phenotypic variance (V*_P_*) is partitioned into variance explained by sequenced SNPs (V_G1_), genetic family structure (V_G2_) and residual variance (V_E_). Final estimates of phenotypic variance partitions are given as proportions. All estimates include standard error.

| Phenotype | $V_{G1}$ | $V_{G2}$ | $V_E$ | $V_P$ | $V_{G1}/V_P$ | $V_{G1+G2}/V_P$ |
| --- | --- | --- | --- | --- | --- | --- |
| Age at first brood (F) | $0 \pm 0$ | $0 \pm 0$ | $0.02 \pm 0$ | $0.02 \pm 0$ | $0.03 \pm 0.19$ | $0.033 \pm 0.1$ |
| Size at first brood (F) | $0 \pm 0$ | $0 \pm 0$ | $0.01 \pm 0$ | $0.01 \pm 0$ | $0 \pm 0.2$ | $0.097 \pm 0.11$ |
| First brood size (F) | $0.78 \pm 1.96$ | $0.12 \pm 2.14$ | $8.86 \pm 1.09$ | $9.76 \pm 0.89$ | $0.08 \pm 0.2$ | $0.092 \pm 0.1$ |
| Interbrood period (F) | $0 \pm 0$ | $0 \pm 0$ | $0.01 \pm 0$ | $0.02 \pm 0$ | $0.17 \pm 0.27$ | $0.302 \pm 0.12$ |
| First brood offspring weight (F) | $0 \pm 0.02$ | $0.03 \pm 0.03$ | $0.06 \pm 0.01$ | $0.09 \pm 0.01$ | $0 \pm 0.28$ | $0.334 \pm 0.12$ |
| Age at maturity (M) | $0 \pm 0$ | $0.004 \pm 0$ | $0.009 \pm 0$ | $0.013 \pm 0$ | $0 \pm 0.2$ | $0.313 \pm 0.09$ |
| Size at maturity (M) | $0 \pm 0.07$ | $0.156 \pm 0.08$ | $0.203 \pm 0.03$ | $0.358 \pm 0.03$ | $0 \pm 0.18$ | $0.434 \pm 0.09$ |

We repeated the analysis on each chromosome to test whether these estimates could be explained disproportionately by certain chromosomes, or whether per-chromosome associations may exist that cannot be observed within genome-wide estimates. Estimates of *h^2^c* based on single chromosome GRMs revealed six chromosomes significantly associated with male size at maturity, four chromosomes with male age at maturity, and one chromosome with offspring weight (FDR ≤ 0.05; Table S7; Figure 2A). Of these however, according to the LRT approach only three chromosomes for male size at maturity (chr5: 20.7%, chr23: 13.9%, chr11: 13.3%), two for male age at maturity (chr1: 14.7%, chr10: 14.4%) and one for offspring weight (chr19: 8.4%) were significantly associated (LRT *p* ≤ 0.05; Table S5). Following multiple-testing correction within phenotypes, only the associations between chr5 and male size at maturity (LRT = 9.346, *fdr* = 0.046), and chr19 and offspring weight (LRT = 9.264, *fdr* = 0.046) were significantly associated according to both methods. Agreement between both methods was good according to a correlation of p-values (Spearman’s ρ = 0.827, *p* < 2.2e−16). Whilst the correlation here is strong, there is a clear downward biasing of p-values from single chromosome GRMs, evident as a shift away from the y=x relationship (Figure S3).

**Figure 2:**
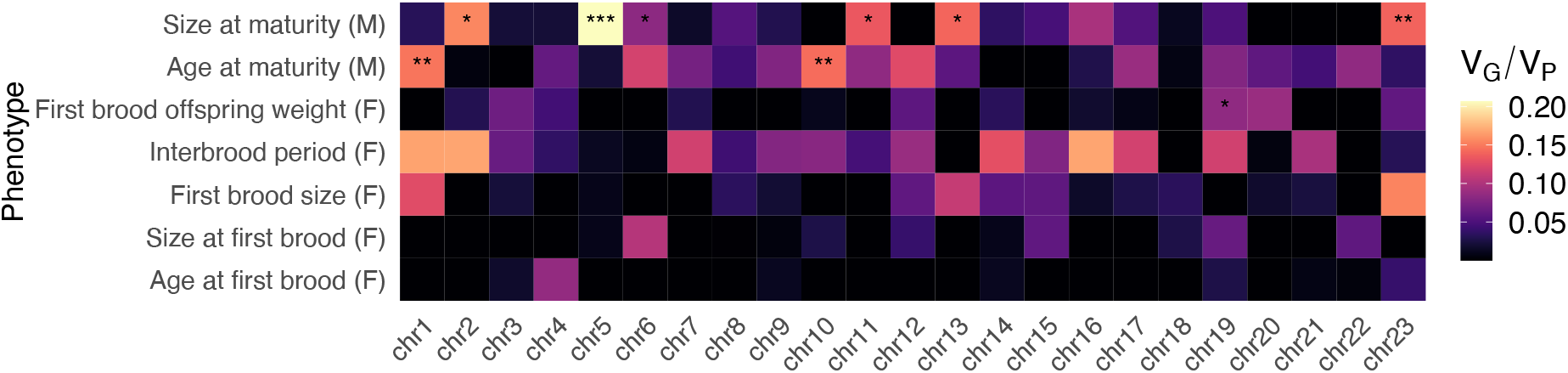
Estimates of phenotypic variance proportions explained by per-chromosome genetic relatedness matrices (*h^2^c*). Tiles are coloured according to the relative proportion of genetic variance (V_G_) to phenotypic variance (V_P_). FDR-corrected p-values (corrected within phenotypes across all chromosomes) are displayed as asterisks FDR ≤ 0.05 = *; FDR ≤ 0.01 = **; FDR ≤ 0.001 = ***).

We found positive correlations associated with single chromosome GRM estimates of *h^2^c* and chromosome size (following the addition of scaffold sizes according to the linkage map; Figure S4) for interbrood period (*r* = 0.373, *p* = 0.04, HC-corrected *p* = 0.105) and male size at maturity (*r* = 0.253, *p* = 0.122, HC-corrected *p* = 231), however these were not significant following HC-correction.

### QTL mapping

We first mapped sex as a binary trait. The location of the sex-determining locus has been narrowed down to a small region at the distal end of chromosome 12 (Fraser *et al.*, 2020; Charlesworth, Bergero, *et al.*, 2020). Our QTL mapping recovered a single large peak on chromosome 12 (17.79 cM), with confidence intervals extending from 5.35-27.78 cM (Figure S5). In our map, the region following this (27.78-61.79 cM) corresponds to the very distal tip of chromosome 12 (approximately 24.6 Mb onwards, plus additionally placed scaffolds), which is the only fully recombining region of this chromosome and is pseudoautosomal (Charlesworth, Zhang, *et al.*, 2020). This places the sex-determining region somewhere in the non-recombining region immediately prior to the pseudoautosomal region (PAR), as proposed by others (Fraser *et al.*, 2020; Charlesworth, Bergero, *et al.*, 2020). This analysis therefore confirms good power to detect loci of large effect and confirms previously published information about the sex chromosome and region containing the sex-determining locus.

We then mapped female traits using all fully informative markers (N = 1220) across all 267 females. Applying a permuted 5% threshold (N=1000), we detected two QTL. The strongest QTL was associated with first brood offspring weight at the very distal tip of chr19 (chr19:66.954) (Figure 3B), explained 7.46% of phenotypic variance, and exhibited additive effects in which the HH homozygotes produced smaller offspring than LL homozygotes (Figure 3C). Confidence intervals (drop in LOD of 1.5) extended between 55.335-67.877 cM. The other QTL was associated with size at first brood (chr22:63.593) (Figure 3D), explained 5.94% of phenotypic variance, and exhibited additive effects in which the HH homozygotes had their first brood at a larger size than LL homozygotes (Figure 3E); contrary to HP-LP expectations. Confidence intervals for the chromosome 22 QTL extended between 44.235 cM – 68.753 cM.

**Figure 3:**
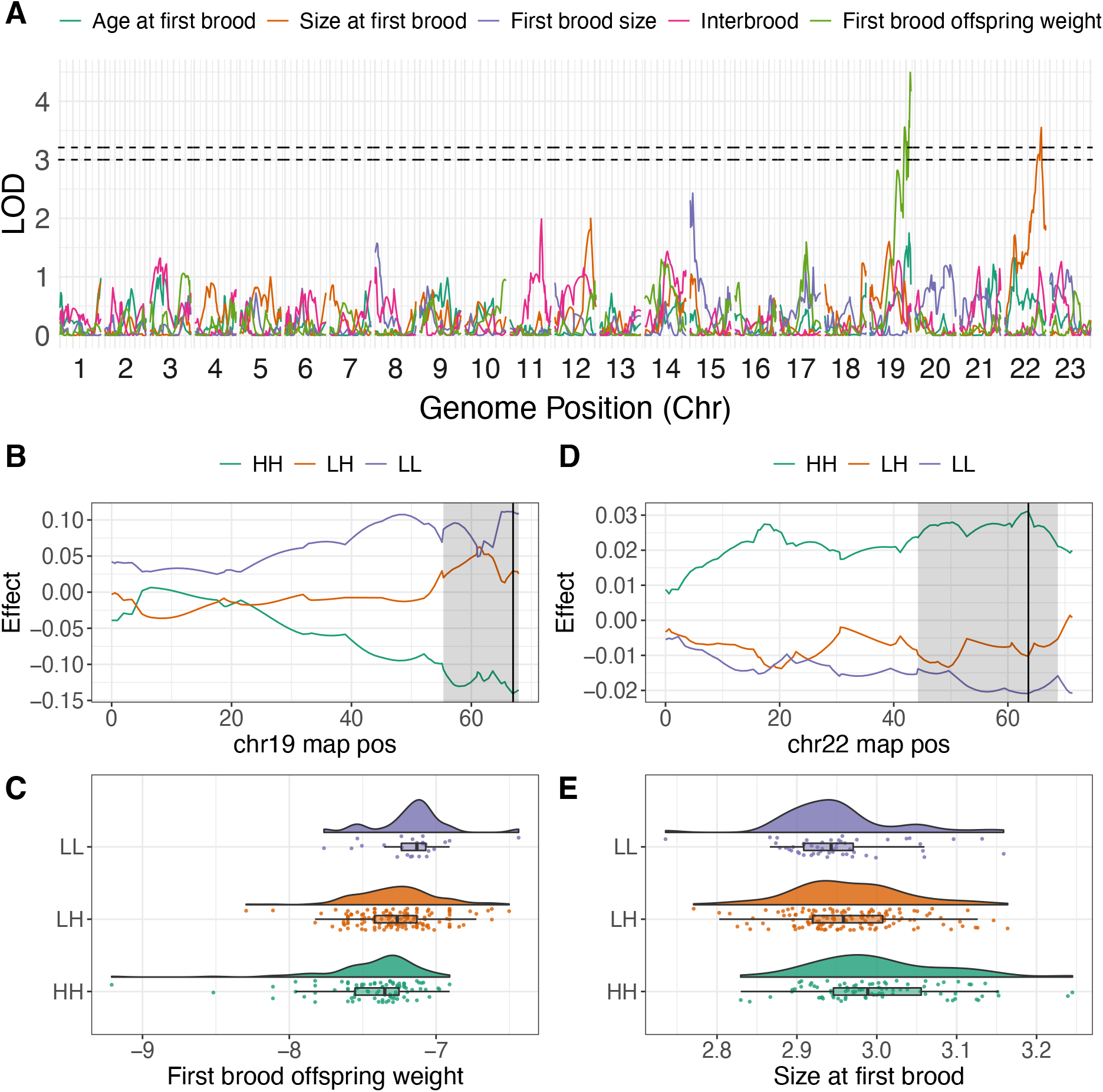
Single locus QTL scans for female life history traits: Age at first brood (days), size at first brood (cm), first brood size (residual), interbrood period (days) and first brood offspring weight (g). All female traits were log-transformed. Panel **A** shows genome-wide additive model scan results for all traits. 5% significance thresholds are denoted. Significant QTLs for offspring weight (chr19, **B**-**C**) and size at first brood (**D**-**E**) are visualised in further detail. Panels **B**-**E** show QTL effects across the focal linkage groups (**B** and **D**), and distributions of phenotypes across genotypes at the peak (**C** and **E**). In panels **B** and **D**, the QTL peak is shown as a black line, with confidence intervals (LOD drop = 1.5) highlighted by grey shaded areas.

Scans within each of the four families identified an additional three QTL (Figure S6-7). Two of these were observed in cross YHF5QM6, associated with first brood size (chr23:37.33, LOD = 3.386, ci low = 4.589 cM, ci high = 53.588 cM) and interbrood period (chr12:60.713, LOD = 3.406, ci low = 54.701 cM, ci high = 72.487 cM). These QTL explained 18.55% and 18.65% of phenotypic variance within their families respectively. The QTL linked with interbrood period here is particularly interesting given its proximity to the sex-determining region. We also detected a QTL associated with female interbrood period in cross QF4YHM8 (chr14:28.41, LOD = 3.559, ci low = 28.412 cM, ci high = 67.316 cM), explaining 17.9% of phenotypic variance for this trait in this family.

We applied the same methodology to male traits, however we did not recover any significant QTLs across the whole dataset (Figure 4). Within the YHF6QM7 cross, we observed a significant QTL associated with male mature length on chromosome 23 (chr23:31.741, LOD = 3.462, ci low = 0 cM, ci high = 52.193 cM) (Figure S8-9). This confidence interval covered the majority of the chromosome, and the QTL explained 9.48% of phenotypic variance in this cross. At this QTL, HP homozygotes matured at a smaller size than LP homozygotes.

**Figure 4:**
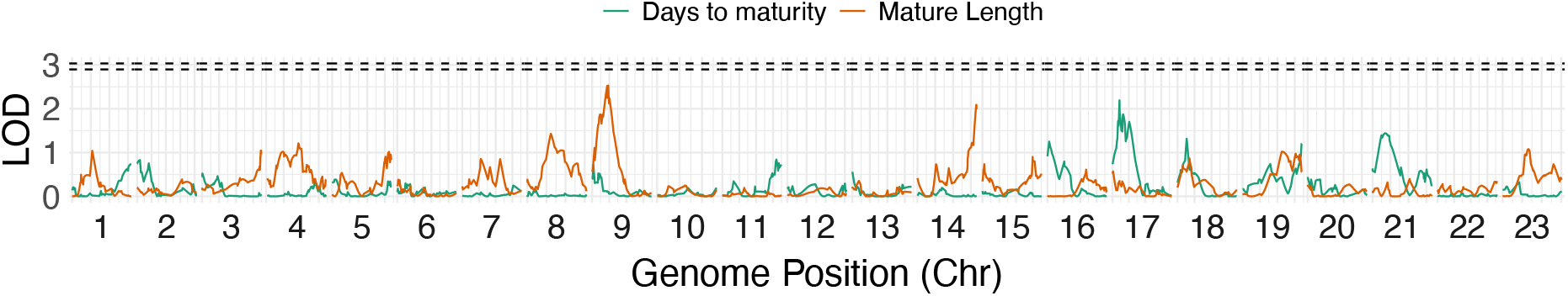
Single locus QTL scans for male life history traits: Days to maturity (log-transformed), length at maturity (mm). No significant QTLs were detected.

We also explored multi-QTL models using r/qtl’s *stepwiseqtl()*, allowing for models that fit up to six loci, with the same additive covariance matrices of family and rearing covariates used previously. For female models, this approach returned null models for all phenotypes except size at first brood and offspring weight, for which we recovered single QTL models including only the QTL loci previously identified above. For males, null models were returned for both phenotypes.

### Candidate genes

Taken together, we find that where traits are heritable, they are associated with whole chromosomes, suggestive of large polygenic regions, rather than single large effect QTL. However, we observed two significant QTL for female traits (offspring weight chr19:66.954 and size at first brood chr22:63.593). We also observe several significant QTL within specific families. Of these, the QTL associated with interbrood in cross YHF5QM6 (YHF5QM6-chr12:60.71) is of particular interest given its proximity to the sex-determining region and relatively narrow confidence intervals. We therefore explored these three regions further for candidate genes. Full information on candidate genes can be found in Table S8.

The QTL at chr19:66.954, associated with offspring weight, covered ~8.2 Mb of chromosome 19, and included 267 genes. Due to the large size of this region, and that alternative confidence intervals (Bayesian 5% probability intervals) suggested a single peak at chr19:66.954, we limited our curation of candidate genes to the immediate 0.5 Mb either side of the peak (chr19:18602889-19602889). This region included 40 genes. Several of these genes (*wfikkn2b*, *tob1b*, *sap30bp*, *h3-3b*, *unk*, *mrpl38*, *fdxr*, *narf*, *cybc1*) are expressed in all stages of embryonic development in zebrafish, or interact with growth signalling pathways, suggesting potential functional effects for offspring weight. The closest gene to the peak was *cdr2l*. Of the 40 genes in this region, 18 exhibited female-biased differential expression in gonads based on the guppy transcriptome (Sharma *et al.*, 2014), and six exhibited male-biased expression (Table S8), suggesting potential reproductive function.

The chr22 QTL peak (for size at first brood) was on a previously unplaced scaffold 000111F_0:651336 and confidence intervals extended over the scaffold (00111F_0:528180-1199617) and a region at the distal end of chromosome 22 (chr22:23415429-24223839). The peak of the QTL did not overlap any genes, however the closest gene was F-box protein 33, *fbxo33*. In total, the region included 45 genes. There were no genes with GO terms or KEGG annotations indicative of clear roles in growth. According to the GeneCards database however, several genes had evidence of affecting size and growth phenotypes in mice knockout studies (*fbxo33*, *tonsl*, *cnr1*, *cga*, *htr1b* and *myo6*). Two of the genes (*cga* and *htr1b*) in this region are also associated with various hormonal pathways, including gonadotropin hormone signalling that may affect development. Other genes in this region are associated with myogenesis, including *myo6* and genes within the Akirin family (*akirin2*, *gabrr1*, *pm20d2*, *cnr1*, *syncripl*). In addition, the gene *snx14* has been associated with growth QTLs in grass carp (Huang *et al.*, 2020).

The interbrood period QTL on chr12 overlapped the YTH domain containing 1 gene, *ythdc1*, on a previously unplaced scaffold 000149F_0 (000149F_0: 131600). This scaffold was placed at the distal end of chromosome 12 near the sex-determining region, and corresponds with scaffold KK215301.1 in the older female genome (Künstner *et al.*, 2016), which has similarly been placed at the distal end of chromosome 12 in other mapping studies (Charlesworth, Bergero, *et al.*, 2020). The confidence interval around this QTL covered additional regions of scaffold 000149F_0 (000149F_0:47124-197258) and chromosome 12 (chr12:24525856-24705290). This region included 18 genes, but none were associated with clear GO or KEGG terms indicative of roles for female fertility. Four genes exhibited female-biased expression in gonads and seven were male-biased. The gene overlapping the QTL peak, *ythdc1*, however is a promising candidate due to its interactions with N6-methyladenosine (*m^6^A*) (Wang *et al.*, 2015; Xia *et al.*, 2018). Disruption of *m^6^A* by mutation of another modifier *mettl3*, affected oocyte development and reduced the proportion of full-growth follicles in zebrafish (Xia *et al.*, 2018). Further, *Ythdc1*-deficient mice have oocyte maturation blocked at the primary follicle stage and experience alternative splicing defects in oocytes (Kasowitz *et al.*, 2018). Examination of transcripts matching *ythdc1* (largest = CUFF_24477_m.316355) in the guppy transcriptome (2014) revealed significant overexpression in ovaries compared with testes. Sequence analysis alongside other Poeciliids (Ensembl release 101: *P. formosa,P. latipinna, P. mexicana,* and *Xiphophorus maculeatus*) demonstrated significant purifying selection on this gene (dN/dS ≤ 0.197; Z-tests for purifying selection: 2.79 ≤ *Z* ≤ 7.40; Table S9). These provide strong evidence for a functional reproductive role for this gene in guppies.

## DISCUSSION

Using an F2 cross design of outbred populations with divergent life history phenotypes, we have demonstrated both polygenic and oligogenic trait architectures underlying guppy life history evolution. For both male size and age at maturity, we find significant heritability associated with particular chromosomes, suggesting polygenic traits, but little evidence of a genome-wide signal of polygenicity or single loci of large effects. For the five females traits, we recovered significant genome-wide estimates of heritability associated with interbrood period and offspring weight. For offspring weight, all per-chromosome estimates of heritability were generally weak, but we did identify a single locus of large effect at the distal end of chromosome 19, suggesting an oligogenic architecture. For interbrood period, we detected a weak genome-wide polygenic signal as an association between chromosome size and per-chromosome heritability and also found evidence of a single locus of large effect on the sex chromosome, LG12, in a single family. Together, these results suggest interbrood period has an oligogenic genetic architecture consisting of a combination of genome-wide polygenic loci and an individual locus of larger effect. We also detected a significant single-locus QTL associated with female size at first brood with a small effect size, but negligible estimates of genome-wide heritability for this trait. Finally, we found no evidence for heritable genetic architectures for female first brood size and age at first brood, although we observed a within-family QTL associated with first brood size on chr23.

The significant genetic components of guppy life history phenotypes found here are in line with previous laboratory rearing estimates of heritability; specifically, the high heritability of male traits, interbrood period, and offspring weight has been documented in laboratory-reared populations under controlled conditions (Reznick, 1982) and in LP-introduction experiments (Reznick *et al.*, 1997). However, there is mixed support for the heritability of female life history traits. Consistent differences between laboratory-reared HP and LP populations for female size and age at maturity and interbrood period have been observed (Reznick, 1982), and similarly, Torres-Dowdall et al. (2012) report consistent differences in female life history traits along a predation gradient in wild-caught vs lab-reared guppies from the Guanapo river. However, estimates of heritability of female age and size at maturity from experimental LP-introductions have shown inconsistent, often negligible, estimates of heritability (Reznick *et al.*, 1997). Interestingly, this latter study postulates that the higher heritability, and more rapid evolution, of male life history traits may involve significant loci associated with the Y chromosome, but we found no evidence here to support this (although see discussion about Y-linked male traits below). Rather, our results of a significant polygenic component of male age and size at maturity also predicts more rapid phenotypic evolution for males, as observed in experimental introductions (Reznick *et al.*, 1997).

Recent work has sought to compare the relative contributions of genetic and plastic effects on guppy life history. HP-LP comparisons are confounded by increased competition for available resources at high densities in LP sites (Reznick *et al.*, 2001), which can result in life history traits that are resource dependent. Across multiple laboratory-reared guppy crosses, Felmy et al. (2021), demonstrated that guppy life histories cluster together in terms of those strongly affected by resource plasticity (predominantly size-related traits), those affected by HP-LP ecotype (including interbrood interval and offspring weight), and those affected by both or neither. For female traits, these results align well with ours, particularly the higher within-family heritability for interbrood period and offspring weight (Table 1), and the absence of strong signatures of genetic architectures associated with female size at maturity. The phenotypic covariances for female traits observed here (Table S6) also agree with the proposed “mosaic” of guppy life history traits (Felmy *et al.*, 2021). Felmy et al. (2021) also demonstrate resource-based plasticity for male age and size at maturity, however these likely operate alongside underlying genetics, in agreement with observations here, and documented in other studies of male guppy life history (Reznick, 1982; Reznick *et al.*, 1997, 2005).

Guppy life history traits are also plastic with respect to other features of local environments. Predator cues, for example, influence female size at maturity (Torres-Dowdall *et al.*, 2012) and growth rate (Handelsman *et al.*, 2013), and resource availability affects female reproductive investment (Reznick and Yang, 1993), the latter contributing comparable phenotypic variance to that associated with heritability. We also found significant associations between rearing conditions for both male age and size at maturity and female age at first brood and interbrood period. These rearing effects reflected small fluctuations in temperature (rearing temperature varied between 23.3-27.1 °C) and date of birth (used as a proxy for other unmeasured rearing conditions), suggesting additional plasticity associated with these phenotypes. Whilst we included rearing effects and family classification as covariates in our models, and controlled for resources during rearing, it is quite possible that additional sources of phenotypic plasticity may have obscured our analyses.

A limitation of our crossing design is that we cannot make across-family comparisons. We controlled for family-specific intercepts in models, either by including kinship information or family status as covariates. However, family status accounted for significant phenotypic variance in five of our seven phenotypes (all except interbrood period and female age at first brood; Table S4). These may be attributed to family-specific alleles segregating across the genome, or Y-specifically, however we cannot separate these within the current dataset. We identified some large effect loci segregating within families (Figure S6-9), however our mapping of male phenotypes in particular will be blind to Y-specific QTL, given all males within a family share the same Y allele. Y-linked loci have been suggested to affect age and size at maturity phenotypes in other poeciliids (Kallman and Borkoski, 1978; Lampert *et al.*, 2010), so there is reason to assume these regions may be important and comprise some of the phenotypic variance associated with family status (which is an upper limit).

Whilst we detected both polygenic and single-locus architectures for life history phenotypes, our study is limited by sample size. In part this is due to the modest brood sizes of guppies, which restricted our ability to generate larger F2 families, and also because the phenotypes considered are sex-specific. It is well known within quantitative genetics studies that small sample sizes can inflate estimates of heritability and QTL effect sizes, a phenomenon termed the “Beavis Effect” (Beavis, 1994; Rockman, 2012; Slate, 2013). In particular, sample size has been demonstrated to inflate p-values when run with single chromosome GRMs with GCTA (Kemppainen and Husby, 2018a), which explains why we recover more modest p-values when comparing single chromosome GRM results to the LRT approach. Our estimates of per-chromosome (*h^2^c*) and genome-wide heritability should therefore be treated with caution. More generally, this combined approach of using single chromosome GRMs and LRTs may be a useful strategy to alleviate issues with modest sample sizes associated with each approach individually. Specifically, this refers to the inflation of significance tests with single chromosome GRMs and issues with model convergence for chromosomes with minimal effects under the LRT approach.

The two main QTL observed here, detected across the whole dataset, reflect only marginal PVE (7.46% for offspring weight on chr19, and 5.94% for female size at first brood on chr22) which is likely inflated by our low sample size. Interestingly, each of these chromosome-phenotype pairings was also detected (marginal significance) by our multi-locus approaches. This suggests that these regions may be reasonably large, such that the peak (from which the PVE is calculated) only represents a portion of the variance explained by the wider region. Neither of these regions have been strongly implicated in HP-LP adaptation before in previous population genomic analyses (Fraser *et al.*, 2015; Whiting *et al.*, 2020), however Whiting et al. (2020) recorded a selection scan outlier within the chr22 QTL interval (chr22:23960000-23970000) in HP-LP comparisons from the Aripo and Madamas rivers. The closest gene to this outlier is *sox11a*. While selection scans of HP-LP comparisons are unable to determine which phenotype selection might be acting on, our results here suggest this region may be involved in female growth.

An absence of prominent large effect loci, as described here for the majority of the traits, would be expected to produce minimal molecular convergent evolution. This is predicted due to redundancy in the mapping of genotype to phenotype, which may allow replicate HP-LP pairs to use different sets of alleles to produce convergent HP-LP phenotypes (Barghi *et al.*, 2020). Limited genomic convergence has been observed in two independent evaluations of natural HP-LP populations (Fraser *et al.*, 2015; Whiting *et al.*, 2020), but appears to be more pervasive in experimental translocations of HP guppies to previously uncolonised LP habitats than naturally colonised LP populations (Fraser *et al.*, 2015). Part of this discrepancy can be explained by the concept of “adaptive architecture”, such that the convergent genetic basis of polygenic traits can be influenced by additional factors such as starting allele frequencies. Starting allele frequencies or amounts of standing variation are likely to be more similar when experimental populations are founded from the same population and/or lack founding bottlenecks compared with naturally-derived HP-LP pairs in different rivers. Empirical evidence for genetic convergence occurring with polygenic traits has been observed for male mating song traits in Hawaiian *Laupala* crickets (Blankers *et al.*, 2019) and myxomatosis resistance in rabbits (Alves *et al.*, 2019), suggesting that genetic architecture alone is not necessarily a constraint on genetic convergence.

Similarly, polygenic traits are predicted to facilitate rapid adaptation. Several experimental studies involving the translocation of guppies into upstream LP habitats (Reznick and Bryga, 1987; Reznick *et al.*, 1997, 2019; Gordon *et al.*, 2009) or manipulating predation within populations (Reznick *et al.*, 1990) have demonstrated that HP-LP adaptive traits evolve rapidly over the course of a few generations. Our findings here, that some of these traits exhibit genetic architectures of many loci of small effect, are in keeping with recent empirical (Barghi *et al.*, 2019) and theoretical work (Bell, 2013; Jain and Stephan, 2017a) suggesting these facilitate rapid adaptation. In this framework, many loci of small effect provide adaptive substrate within populations to rapidly respond to shifting optima. This is particularly true if distance to new fitness optima is short, such that environmental change is modest and fitness effects are relative and/or non-lethal. This may be the case for guppy life history traits under LP regimes, where soft selection is most likely. This model therefore could allow male life history traits to change rapidly through small changes at many loci, whilst additional segregating larger effect loci may act in concert with compensatory changes at small effect loci for female traits. Our results here provide regions of the genome and candidate genes to explore further. For instance, an appreciation that much of the genetic basis of guppy life history traits may be polygenic informs on experimental and sampling designs for future population genetic studies of this system. In particular, temporal sampling and quantifying genome-wide autocovariances of neutral allele frequencies offers a promising avenue for studying the role of polygenic architectures in rapid adaptation (Buffalo and Coop, 2019). The genomic regions identified here may serve as focal regions in these studies.

In conclusion, we used an F2 cross to explore the genetic architecture of seven guppy life history traits that are known to evolve rapidly and convergently in natural populations. We find evidence of only two loci of large effect associated with female size at first brood and offspring weight, and evidence of many loci of small effect associated with male age and size at maturity, interbrood period and brood size. In addition, we observed several within-family loci of large effect, suggesting segregating variation within source populations. These results have important implications for improving our understanding of how life history traits evolve in the guppy model, and more broadly, provide empirical evidence for predictions of the genetic architecture of rapidly adapting and convergent phenotypes.

## ACKNOWLEDGEMENTS

The authors wish to thank all members of the Fraser group for useful discussions. Pasi Rastas provided advice for linkage mapping with Lep-MAP3. Simon Zhu, Blanca Guzman, Sara Ruckman, Ruchittrani Hapuarachchi, Christopher Tan, Kevin Khuu, Vicent Poon, Carol Villacana, and Brianna Paramo assisted in fish rearing and crossing at UC Riverside. HPC infrastructure support was provided by The University of Exeter’s High Performance Computing (HPC) facility (ISCA). DNA sequencing was performed by University of Exeter Sequencing Service (ESS).

## DATA AVAILABILITY

All sequencing read data is available from the ENA (DOI: XXX)

All scripts and other data associated with analysis will be made available in an archived github repository (Zenodo, DOI: XXX)

