## Supplementary material for "On the genetic architecture of rapidly adapting and convergent life history traits in guppies": Figure S

**Supplementary materials**

**Author list:**

James R Whiting^1^, Josephine R Paris^1^, Paul J Parsons^1,2^, Sophie Matthews^1^, Yuridia Reynoso^4^, Kimberly A. Hughes^3^, David Reznick^4^, Bonnie A Fraser^1^

**Contents:**

- Supplementary Methods
- Figure S1 - Phenotype distributions by cross family
- Figure S2 - Linkage map
- Figure S3 - Association between single GRM and LRT approach p-values
- Figure S4 - Correlations between *h^2^c* and chromosome size
- Figure S5 - QTL scan results for sex
- Figure S6 - Within-family single-locus QTL scans for female life history traits
- Figure S7 - QTL effect plots for within-family female traits
- Figure S8 - Within-family single-locus QTL scans for male life history traits
- Figure S9 - QTL effect plots for within-family male traits
- Table S6 - PCA results for female life history traits
- Table S9 - dN/dS results across Poeciliids for *ythdc1* candidate

Note: Tables S1-5,7-8 are available as external .xlsx files

**SUPPLEMENTARY METHODS:**

***Phenotyping and Rearing***

Life history phenotyping and rearing followed (Reznick, 1982); full rearing details are available in the supplementary materials. In brief, fish were reared from birth to 25 days in groups of 5 or 6 in two gallon tanks, then sexed and isolated. They were then reared one per aquarium on quantified rations with opaque barriers between tanks. At first, males were checked weekly to evaluate the development of the anal fin, which was used as the index of maturity. They were then checked as often as daily as the fin approached full metamorphosis, as judged from the development of the apical hook and the extension of the hood, with maturity decided when the hood reached or extended beyond the tip of the fin. Females were mated weekly, beginning shortly after they entered the phase of quantified rations. We scored their age when they first gave birth. From then on, females were only mated the day after parturition when they are receptive to mating. All offspring were preserved on the day of birth and the female was preserved after the birth of the third litter. Tanks were cleaned every other week. Size of females and males was measured after they had been anaesthetized with MS-222, with length being measured under a dissecting scope with Vernier calipers. Based on the allometric dependency of female brood size, we looked at the residual brood size derived from comparing the observed brood size with the linear-predicted brood size based on female size at maturity. Where necessary, phenotypes were log-transformed to improve fit for normality assumptions.

***Genotyping***

Genomic DNA was extracted from fin clips using an ammonium acetate extraction method (Nicholls *et al.*, 2000; Richardson *et al.*, 2001). We genotyped each individual using a RAD-seq library preparation method adapted from Poland and colleagues (Miller *et al.*, 2007; Baird *et al.*, 2008; Poland *et al.*, 2012). Briefly, double-digested genomic DNA (enzymes: *PstI* and *MseI*) was annealed with cut-site specific sequencing adaptors bearing individual barcodes. Barcoded samples were amplified via PCR (12 cycles, with barcoded samples multiplexed). Each individual had a unique barcode, using the original barcodes from Poland et al. (2012). A total of 12 multiplexed libraries were sequenced. Of the 661 individuals used in the final analysis (including eight grandparents), 61 individuals were sequenced two or three times in separate libraries to account for low coverage (“merged” individuals in Table S2). To ensure optimal coverage of the grandparents, and to reduce the effects of PCR duplicates, each of the eight grandparents were sequenced four times in four separate PCR reactions and sequencing libraries. Of the total 653 F2s, 637 (370 males, 267 females) were used for phenotype analyses due to missing phenotype for 16 individuals.

***Heterochiasmy and linkage mapping***

There were some differences between male and female maps prior to averaging on some linkage groups, which is expected given female-biased heterochiasmy (Bergero *et al.*, 2019), however these differences were minor and generally reflected differences of intercept rather than slope when comparing genetic maps to the physical genome. These sex-averaged maps therefore are accurate representations of the relative genetic map, but are limited in terms of inferring accurate estimates of recombination rate.

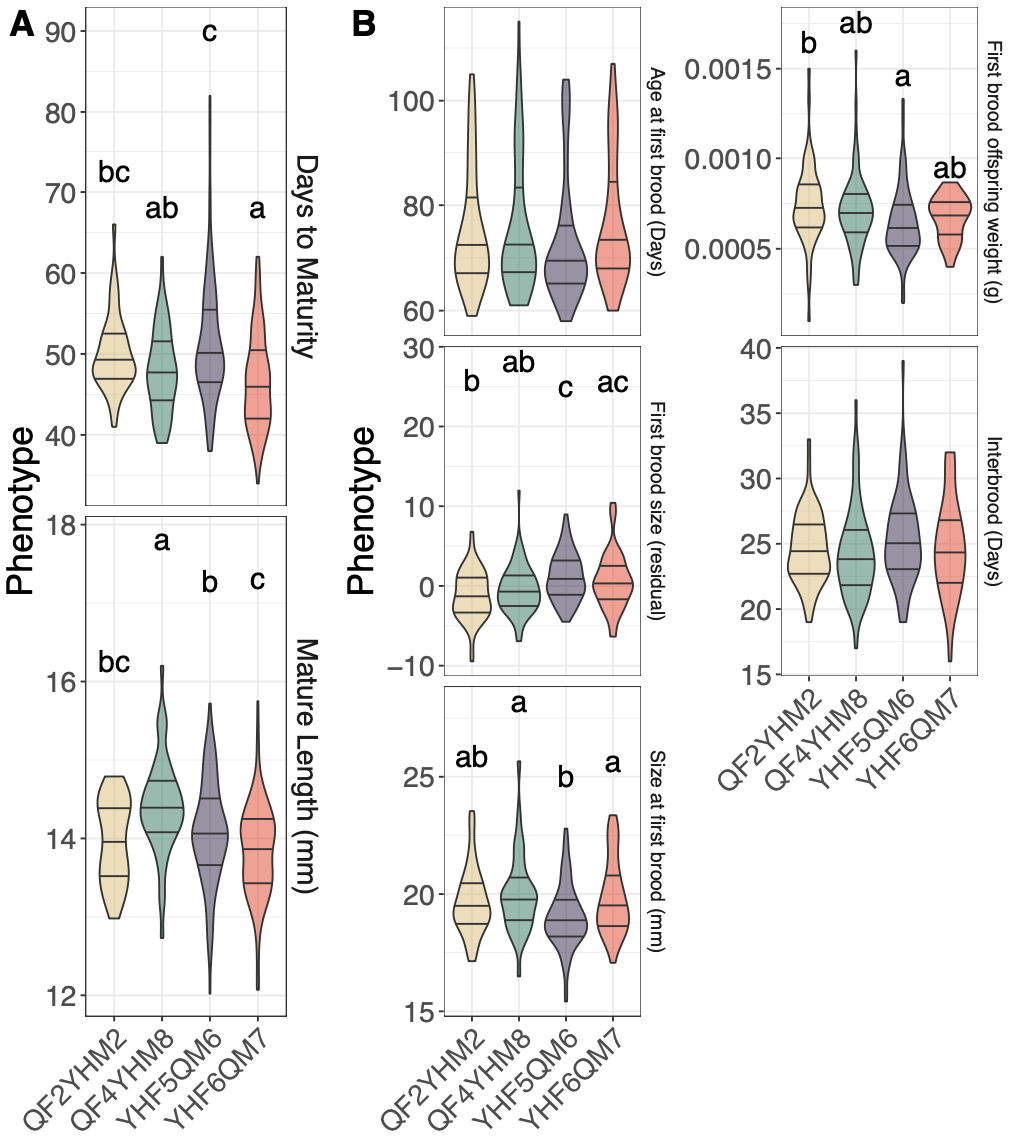

**Figure S1:** Phenotype distributions for male (**A**) and female (**B**) life history phenotypes grouped between families. Where phenotypes differed significantly between families, significance groups are highlighted above violins. Each violin shows the median and quartiles.

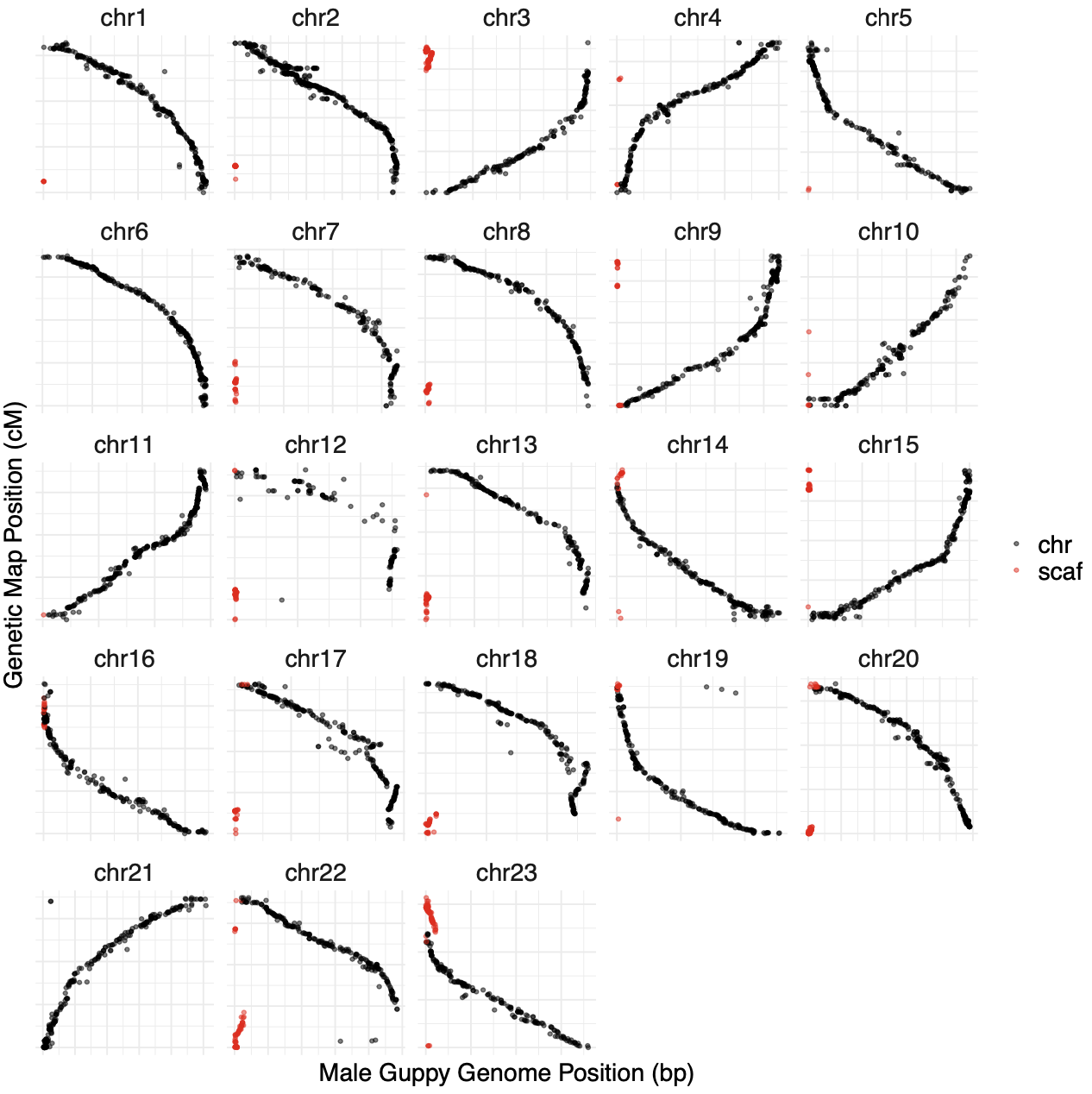

**Figure S2**: Comparison of genetic map positions against male guppy genome. Each facet highlights one of the 23 chromosomes in the genome. Black points represent markers on scaffolds previously mapped to chromosomes, markers in red highlight additional chromosomal regions made up of markers from unplaced scaffolds (Table S2). Positions on the x-axis represent genome positions in bp, always starting at 0. Y coordinates reflect genetic map position, which in some cases is reversed relative to the published genome. Genetic map positions were reversed to match published genome order for final estimates of QTL loci and whole-genome visualisations of QTL LOD distributions.

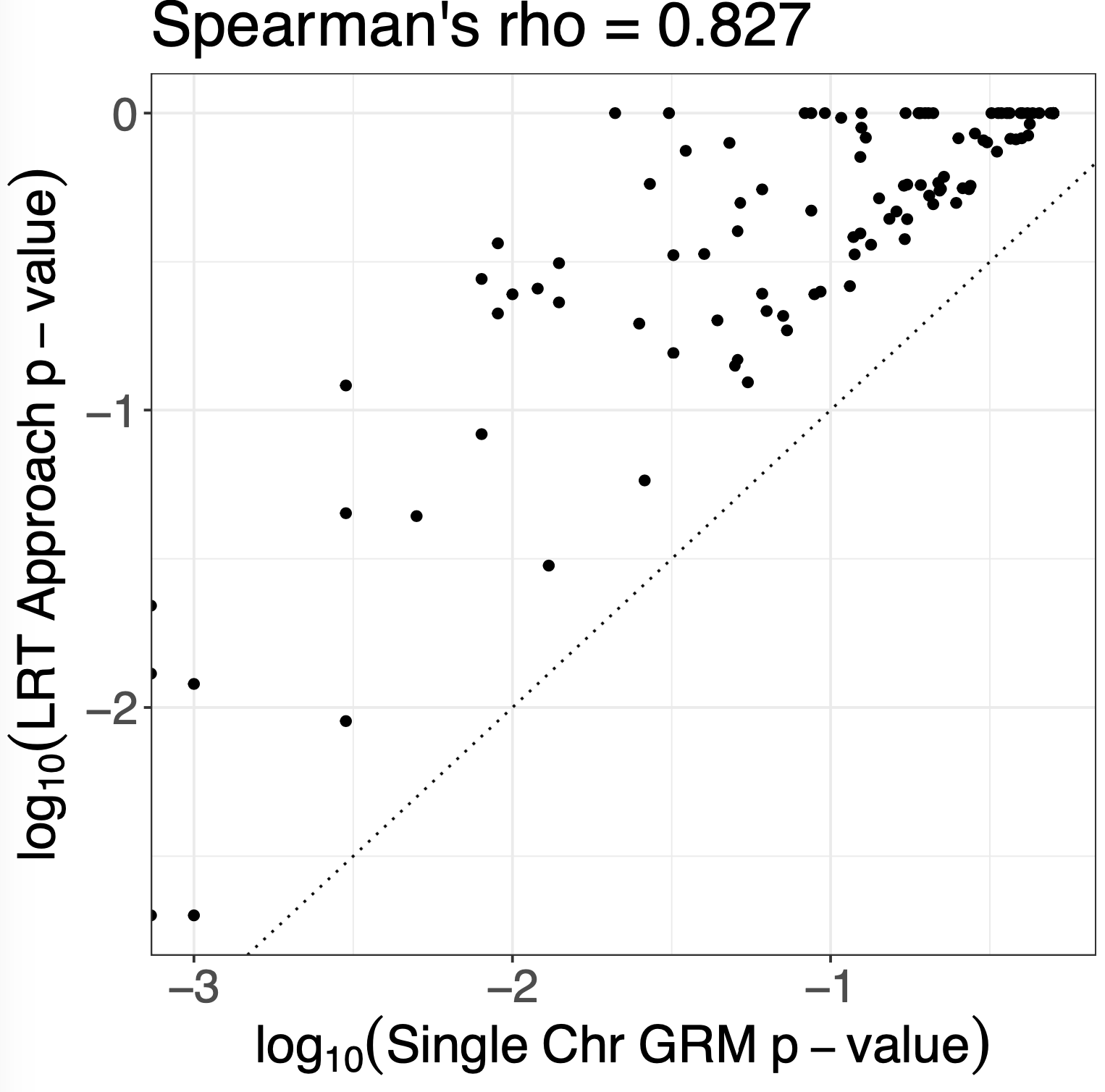

**Figure S3**: Linear relationship between log10-transformed p-values derived from either single chromosome GRM and LRT-approach estimates of *h^2^c*. The y=x relationship is shown as a dotted line. Downward-biasing of single chromosome GRM p-values is evidenced as a shift off the y=x.

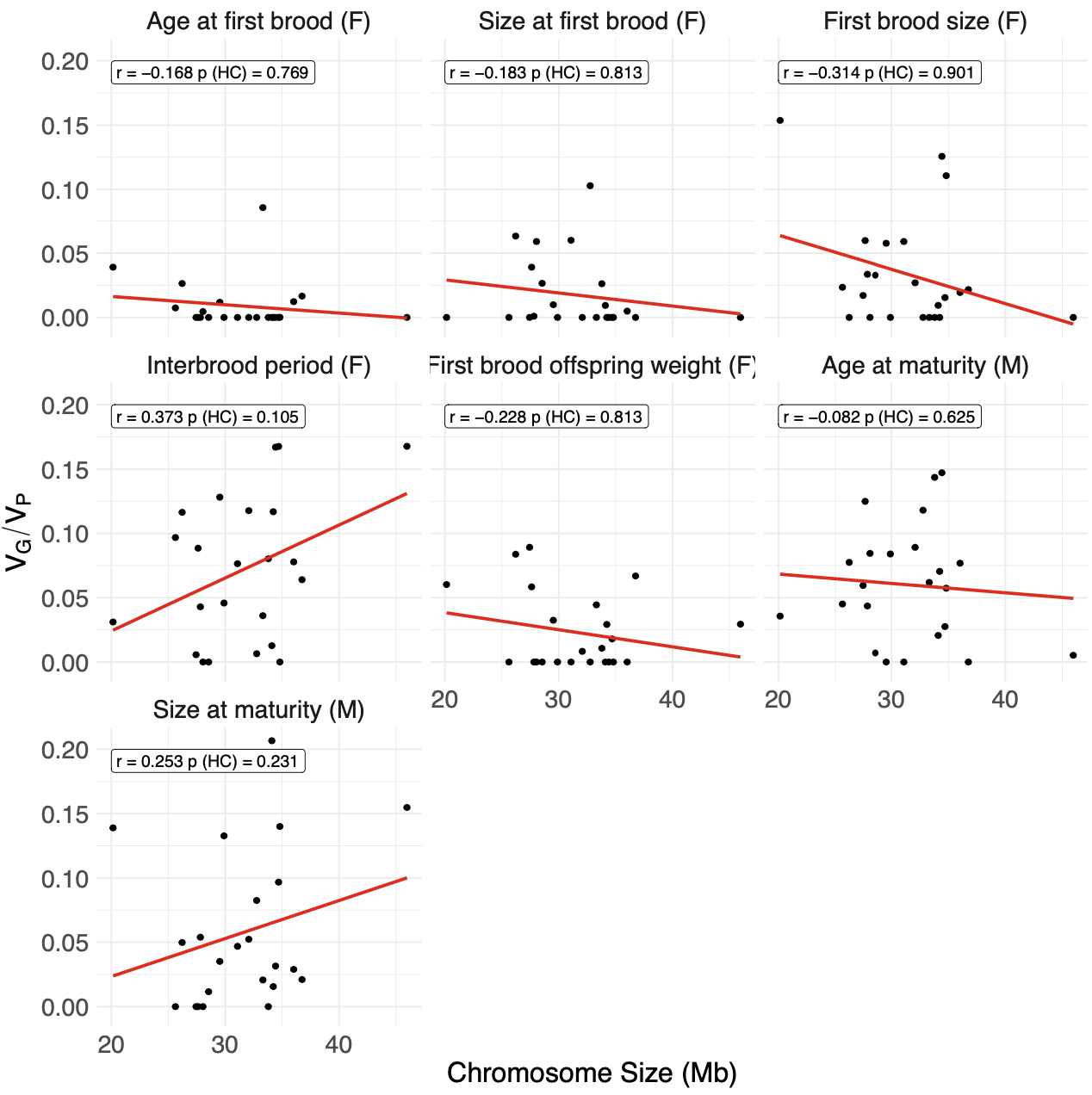

**Figure S4**: Correlations between single chromosome GRM h^2^c estimates and chromosome size. Chromosome size was estimated at the cumulative size of chromosomes and any scaffolds that could be merged to chromosomes according to the linkage map (Figure S2). HC-correction was performed following Kemppainen and Husby (2018), and HC-corrected p-values are shown alongside Pearson’s correlation coefficients within each facet.

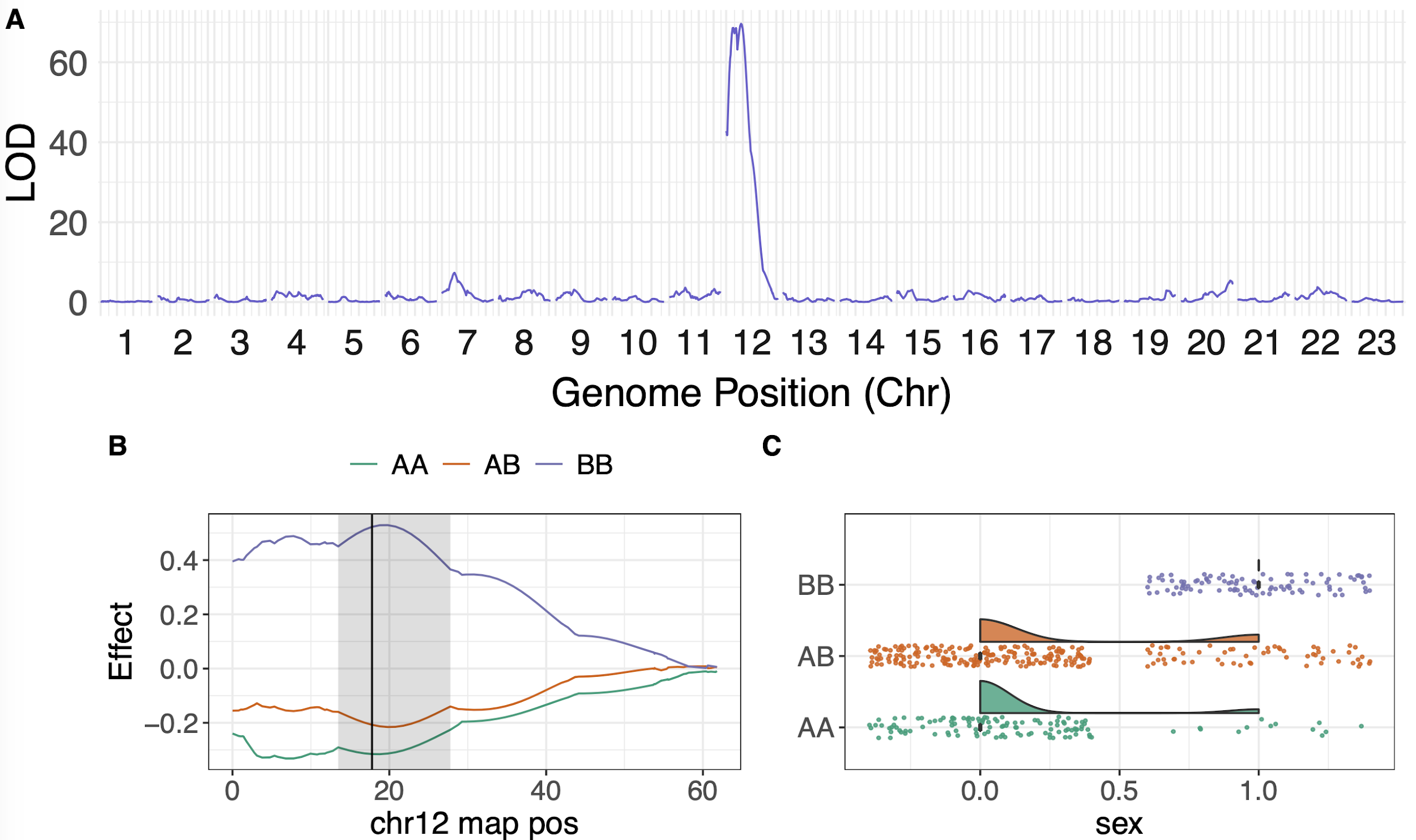

**Figure S5**: Single-locus QTL scan for binary sex classification across all males and females. Models were performed on a binary trait, where male = 0 and female = 1. For this model, genotypes were labelled such that “A” genotypes were inherited from the grandfather, and “B” genotypes were inherited from the grandmother, regardless of cross direction. Panel **B** shows genotype effects across LG12, with the peak denoted as a black line with grey shaded regions showing confidence intervals (drop LOD = 1.5). Panel **C** shows distributions of sex at the peak between the three genotypes, and highlights the expected distribution of a Y-linked marker, such that males cannot inherit both B alleles from the grandmother.

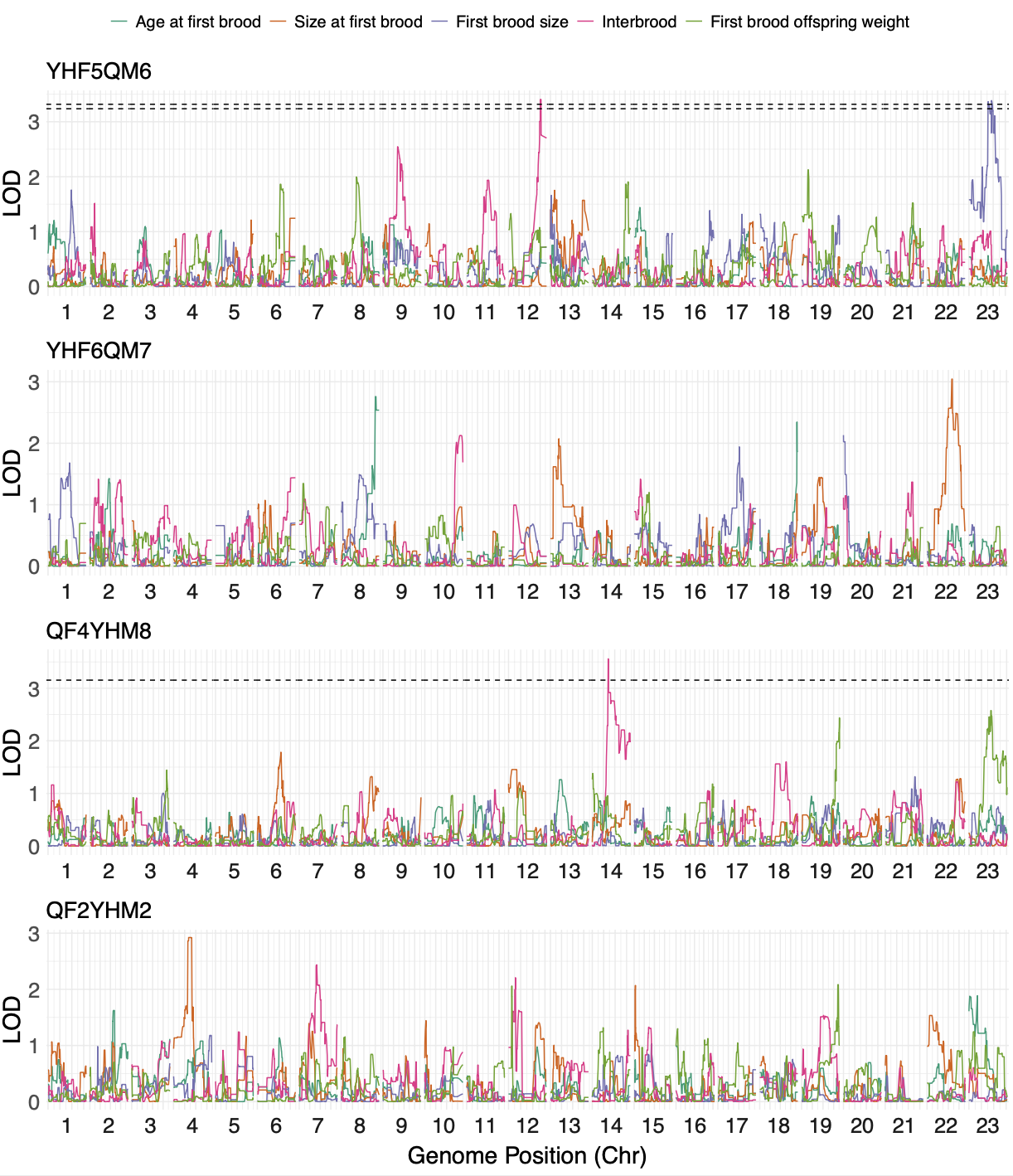

**Figure S6**: Within-family single-locus QTL scans for all five female life history traits. Each row highlights LOD scores across the genome for each of the four crosses. Where significant QTL were detected, 5% permuted (N = 1000) significance thresholds are shown as dashed lines. This analysis highlighted three within-family QTL: two in YHF5QM6, and one in QF4YHM8.

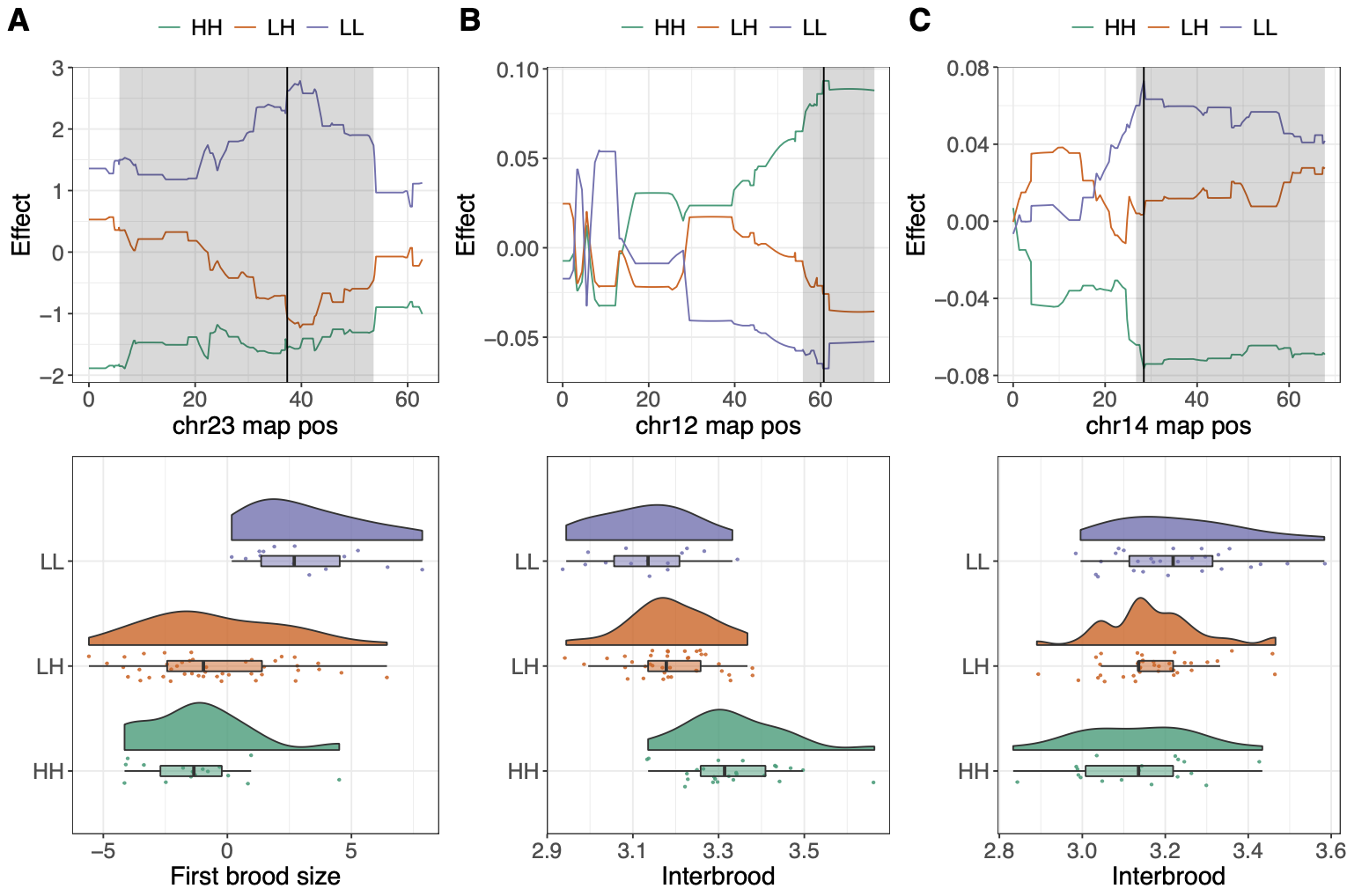

**Figure S7**: QTL effect plots for female within-family QTL detected in families YHF5QM6 (**A**-**B**) and QF4YHM8 (**C**)**.** Each column shows QTL effects across the focal linkage groups (first row), and distributions of phenotypes across genotypes at the peak (second row). For focal chromosomes, the QTL peak is shown as a black line, with confidence intervals (LOD drop = 1.5) highlighted by grey shaded areas. QTL were associated with first brood size (**A**) and female interbrood period (**B**-**C**)**.**

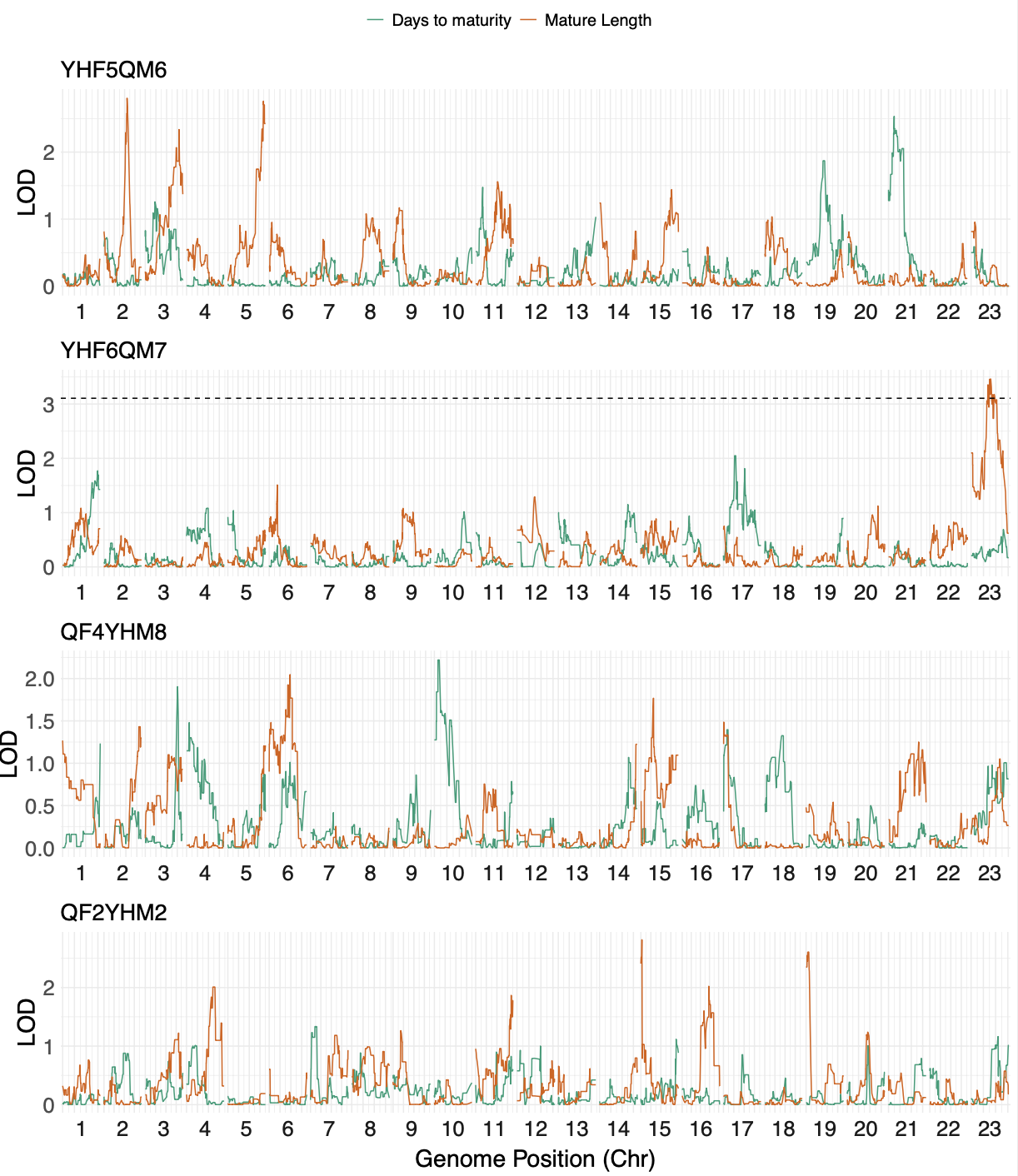

**Figure S8**: Within-family single-locus QTL scans for the two male life history traits. Each row highlights LOD scores across the genome for each of the four crosses. Where significant QTL were detected, 5% permuted (N = 1000) significance thresholds are shown as dashed lines. This analysis highlighted one within-family QTL in YHF6QM7.

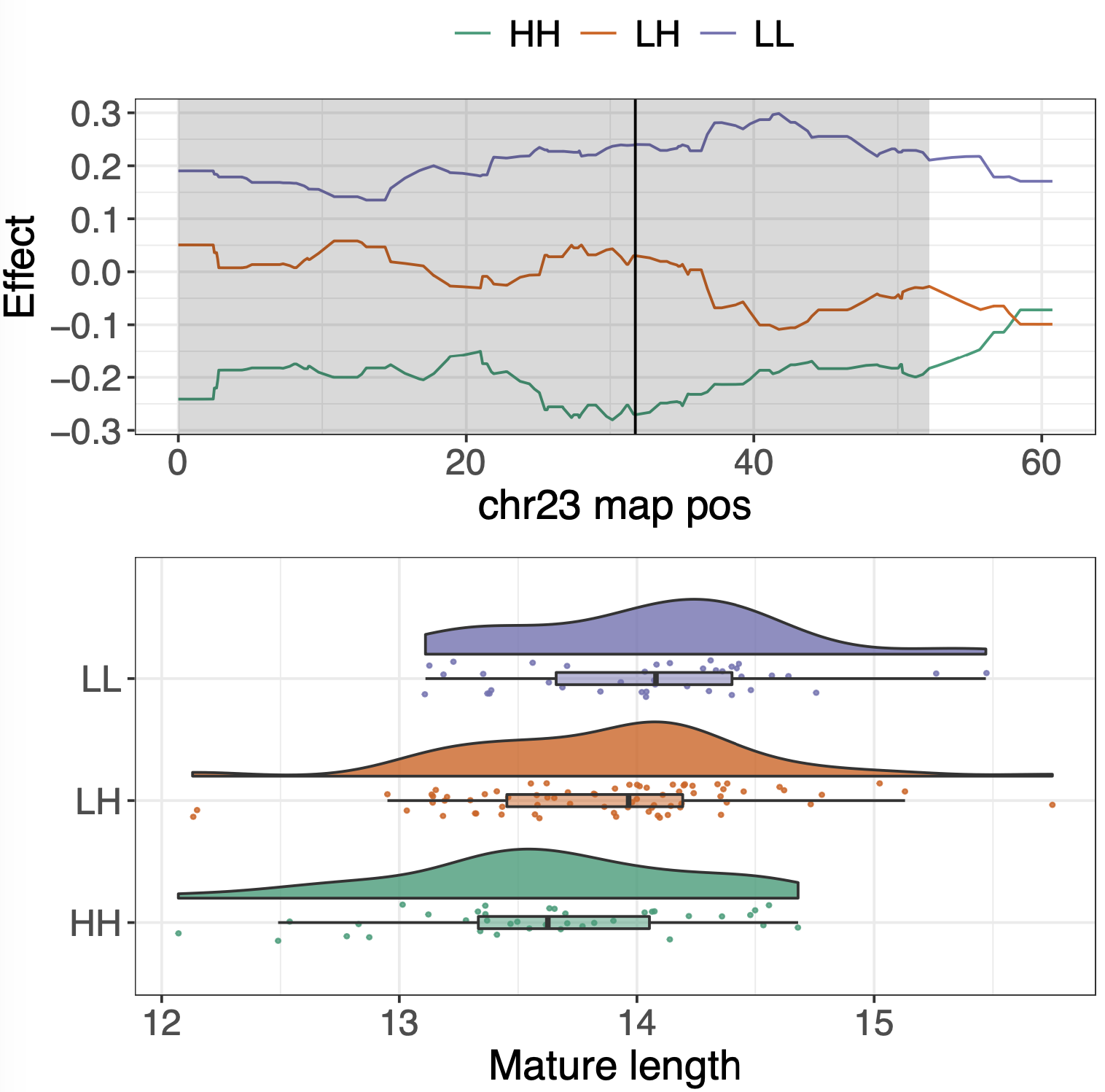

**Figure S9**: QTL effect plot for the male within-family QTL detected in family YHF6QM7**.** The first row shows QTL effects across the focal linkage groups, and the second row shows the distributions of phenotypes across genotypes at the peak. For the focal chromosome, the QTL peak is shown as a black line, with confidence intervals (LOD drop = 1.5) highlighted by grey shaded areas. This QTL was associated with male length at maturity.

| **Table S6:** Principal component analysis between female phenotypes, highlighting positive covariance among age and size at first brood and first brood size. | | | | | |
| --- | --- | --- | --- | --- | --- |
| Phenotype | PC1: 37.6% | PC2: 27% | PC3: 17.7% | PC4: 13.4% | PC5: 4.3% |
| Age at first brood | 0.684 | -0.023 | -0.052 | -0.086 | -0.722 |
| Size at first brood | 0.58 | 0.278 | -0.399 | -0.26 | 0.6 |
| First brood size (residual) | 0.251 | -0.618 | -0.222 | 0.684 | 0.192 |
| Interbrood period | 0.208 | -0.587 | 0.565 | -0.489 | 0.234 |
| Offspring weight | 0.299 | 0.443 | 0.686 | 0.467 | 0.164 |

| **Table S9**: dN/dS values associated with the *ythdc1* gene across the guppy and its close relatives. | | | | | |
| --- | --- | --- | --- | --- | --- |
|  | P reticulata | P formosa | P latipinna | P mexicana | Xiphophorus maculatus |
| P reticulata | 0 | 0.167 | 0.19 | 0.194 | 0.197 |
| P formosa | 0.167 | 0 | 0.11 | 0.114 | 0.12 |
| P latipinna | 0.19 | 0.11 | 0 | 0.111 | 0.148 |
| P mexicana | 0.194 | 0.114 | 0.111 | 0 | 0.111 |
| Xiphophorus maculatus | 0.197 | 0.12 | 0.148 | 0.111 | 0 |
